# Performance of Turquoise killifish, model organism in aging, on commercial pelleted diet: a step towards husbandry standardization

**DOI:** 10.1101/770479

**Authors:** Jakub Žák, Iva Dyková, Marin Reichard

## Abstract

Dietary alteration is one of the most universally effective aging interventions, making its standardization a fundamental need for model organisms in aging. Here we address the current lack of standardized formulated diet for Turquoise Killifish *Nothobranchius furzeri* – a promising model organism. We first demonstrated that *N. furzeri* can be fully weaned onto a standardized commercially available pelleted diet as the sole nutrition when kept in social tanks. We then compared nine somatic and six reproductive parameters between fish fed a typical laboratory diet - frozen chironomid larvae (bloodworms) and fish fed solely on BioMar pellets. Killifish readily consumed the pellets. Although fish consumed 7.5 times less food mass in the form of pellets than bloodworms, they had comparable somatic and reproductive performance. There was no difference between diet groups in body size, specific growth rate, condition or extent of hepatocellular vacuolation. Fish fed a pelleted diet had higher juvenile body mass and more visceral fat. Pellet-fed males had lower liver mass and possessed a lipid type of hepatocellular vacuolation instead of the prevailing glycogen-like vacuolation in the bloodworm-fed group. No significant effect was found on reproductive parameters. The negligible differences between dietary groups and good acceptance of pellets indicates their suitability as a useful starting point for diet standardization (and potential manipulation) in *Nothobranchius furzeri*.

## INTRODUCTION

A standardized diet is an important prerequisite in studies of the mechanisms underlying the biological phenomenon of aging (Nakagawa et al. 2012). Hence there is a high demand for the standardization of laboratory diets (Polačik et al. 2016; Dodzian et al. 2018). Feeding laboratory organisms live food of wild origin has numerous drawbacks such as the risk of disease introduction (Watanabe et al. 1991), chemical contamination of food affecting fish physiology (Papa et al. 2019), seasonal availability or instability of nutritional content (Armitage 1995) and high waste production (Watanabe et al. 1991). All these problems can be avoided by a standardized pelleted diet. Moreover, experimental studies benefit from easy manipulation of the nutritional content of the formulated diet and its effect on aging, lifespan, growth or reproduction (Spitsbergen et al. 2012; Moatt et al. 2019). A standardized diet is often not available for newly introduced laboratory animals. For example, the short-lived Turquoise killifish *Nothobranchus furzeri* is a relatively new model organism in biomedicine and evolutionary ecology (Hu & Brunet 2018; Cellerino et al. 2016; Genade et al. 2005) and no standardized diet is yet available for this species (Polačik et al. 2016; Dodzian et al. 2018); *Nothobranchius* spp are reluctant to accept dry food (Polačik et al. 2016). The absence of a standardized diet likely impedes wider use of Turquoise killifish as a laboratory model (Reichard & Polačik 2019).

Diet has a profound effect on survival and aging of experimental animals and it is one of the key components in aging intervention studies (Moatt et al. 2019; Nakagawa et al. 2012). Diet affects lifespan probably via a maintenance-reproduction trade-off (Shanley & Kirkwood 2000) and hence there is a need to understand the effect of diet on a wide range of life histories. Indeed, the type of diet has different effects on life histories in many fish species. For example in zebrafish *Danio rerio* and Siamese fighting fish *Betta splendens*, growth, condition, fecundity and fertilization rate vary between live food and a pelleted diet (Markovich et al. 2007; Mandal et al. 2012). The impact of diet on life history outcomes thus presents an excellent measure for testing the effect of a pelleted diet on *N. furzeri* compared to the commonly used frozen bloodworms (*Chironomous* larvae) (Polačik et al. 2016).

Diet usually influences energetic reserves stored in livers (Wolf & Wheeler 2018). Energetic reserves in the form of glycogen or lipids refer to hepatocellular vacuolation (Wolf & Wheeler 2018). Fish hepatocytes tend to be more vacuolized than mammal hepatocytes (Wolf & Wolfe 2005) which are often misdiagnosed as lipidosis or steatosis in fish (Wolf et al. 2015). Findings from wild annual killifish suggest that high hepatocellular vacuolation is a natural phenomenon (Vrtílek, Žák, Blažek, et al. 2018; Godoy et al. 2019) which is understandable for a species with high energy demands on growth and reproduction in the unpredictable environment of ephemeral pools (Reichard & Polačik 2019). There is no strong empirical evidence that highly vacuolated livers with compact cellular membranes are not functioning properly (Wolf et al. 2015), though the degree of vacuolation when the hepatocellular membrane disintegrates and neighboring cells fuse is suggested to be pathological condition in the majority of fish species (Wolf & Wolfe 2005). Liver histopathology is one of the most important markers in *Nothobranchius* aging studies (Di Cicco et al. 2011; Terzibasi Tozzini et al. 2009) and it is necessary to provide baseline data for the introduction of a new diet.

*Nothobranchius furzeri* is an important vertebrate model in biomedical and evolutionary studies on aging (Reichard & Polačik 2019; Hu & Brunet 2018; Genade et al. 2005). It has an unprecedented fast life history adapted to shallow ephemeral savanna pools in southern Mozambique and Zimbabwe (Reichard & Polačik 2019). It is a short-lived vertebrate (Valdesalici & Cellerino 2003), with a lifespan of a 1-5 months in the wild (Vrtílek, Žák, Polačik, et al. 2018) and 3-16 months in captivity (Valdesalici & Cellerino 2003; Polačik et al. 2014). In the wild, fish hatch from eggs buried in the sediment after rain-fed pools are flooded and reach sexual maturity within two weeks when they can measure up to 55 mm (Vrtílek, Žák, Pšenička, et al. 2018). They spawn daily and females lay usually 20 - 120 eggs each day (Vrtílek & Reichard 2016). When the ephemeral pool desiccates, the adults die and only diapausing eggs in the pool substrate survive the dry period (Reichard & Polačik 2019). Their natural diet consists of small aquatic invertebrates (Polačik & Reichard 2010). In the laboratory, dietary restriction (Terzibasi Tozzini et al. 2009) and supplementation with antioxidant resveratrol (Valenzano et al. 2006) has been shown to extend the lifespan of *N. furzeri*, demonstrating the significant role of dietary manipulations. Unfortunately, the absence of a standardized diet limits the validation of experimental results across different laboratories (Polačik et al. 2016; Dodzian et al. 2018).

At present, the typical laboratory diet fed to *Nothobranchius* spp. consists of frozen bloodworms (larvae of Chironomidae) but there is considerable variation in the quality of bloodworms from different commercial suppliers and consequently among different laboratories (Polačik et al. 2016; Dodzian et al. 2018). Less common forms of diet used for adult nothobranchids include gelatin cubes made of bloodworms (Valenzano et al. 2006), freeze-dried bloodworms (Almaida-Pagán et al. 2018; Valenzano et al. 2011), *Tubifex* sp. worms (Blažek et al. 2013), *Chaoborus* larvae (Genade et al. 2005) or a combination of *Artemia nauplii* and flake food (Hsu & Chiu 2009). Undoubtedly, the above mentioned studies would have benefited from the existence of a standardized diet. There have been attempts to develop a standardized diet for *N. furzeri* (Powell et al. 2016) but to our knowledge, unfortunately, full conversion to standardized diet has never been successful and co-feeding with live *Artemia* is necessary. *Nothobranchius* spp. strongly prefer natural food and usually avoid the common, commercial dry or gelatin-like fish foods accepted by many other fish models.

In the present study, we used a wild-derived strain of *N. furzeri* (MZCS 222) kept in social groups to investigate their ability to accept the commercially available dry pellets BioMar INICIO. This formula was developed to meet the nutritional requirements of juvenile salmonids, the natural diet of which resemble that of nothobranchids. After successful conversion of fish to dry pellets, we compared key fitness-related life history traits between the pellet-fed diet group and a bloodworm-fed control. We compared growth, Fulton’s condition factor, visceral fat score, liver mass, liver histology, fecundity, fertilization rate, reproductive allotment, egg size, egg survival and egg development dynamics between the two dietary groups. We believe that the present study makes a significant contribution to the standardization of a laboratory diet for this important model organism.

## RESULTS

All fish selected for the pellet diet treatment were successfully weaned from *Artemia* nauplii and bloodworms to pellets between the age of 12 and 21 days, with most fish weaned after five days (age 17 days). The macronutrient content of the diets is provided in Table 1; pellet food had a higher lipid content (11-22% vs. 5%) while bloodworms were richer in carbohydrates (23% vs 6-9%).

**Table 1:** Macronutrient composition of diets fed to Turquoise killifish.

|  | <sup>a</sup><br>INICIO<br>0.4 mm | <sup>a</sup> INICIO<br>0.6 mm | <sup>a</sup> INICIO<br>0.8 mm | <sup>a</sup> INICIO<br>1 mm | <sup>a</sup> INICIO<br>1.5 mm | <sup>b</sup><br>Bloodworms | <sup>b</sup><br>INICIO 0.8 mm |
| --- | --- | --- | --- | --- | --- | --- | --- |
| Crude protein | 63.0 % | 62.0 % | 56.0 % | 56.0 % | 54.0 % | 53.8 % ± 3.0 % | 59.5 % ± 3.0 % |
| Crude lipid | 11.0 % | 13.0 % | 18.0 % | 18.0 % | 22.0 % | 5.2 % ± 8.0 % | 17.2 % ± 8.0 % |
| Carbohydrates<br>(NFE) | 7.3 % | 6.3 % | 9.0 % | 9.0 % | 7.7 % | 22.8 % ± 20.0 % | 11.0 % ± 20.0 % |
| Ash | 12.5 % | 12.5 % | 11.5 % | 11.5 % | 11.0 % | 18.1 % ± 2.5 % | 12.3 % ± 2.5 % |
| Moisture | - | - | - | - | - | 82.9 % ± 2.0 % | 5.9 % ± 2.0 % |

<sup>a</sup> Official declaration of the manufacturer - BioMar INICIO, Denmark
<sup>b</sup> Values from our analysis. Percentages of Crude protein, Crude lipid, Carbohydrates and Ash are computed after subtraction of moisture. Values behind ± are precision of the analytical measurements. NFE stands for nitrogen free extract.

**Table 2:** Overview of statistical models with sample sizes. The number behind the slash symbol is total number of eggs analysed. Models with missing sex factor were performed separately for each sex due to significant sex-specific interaction (or only completed for females in the case of reproductive parameters). LME – Linear mixed effect model (Bates et al. 2015); GLMM – Generalized linear mixed effect model (Bates et al. 2015). In all binomial models raw data were used instead of percentages to account for sample size. MPA – Multistratum permutation analysis with specified 10000 iterations (Wheeler & Torchiano 2016). χ2 analyses were completed by Pearson’s chi-squared test with simulated p-value (based on 10000 replicates). J: juvenile; M: male; F: female. For parameter coding refer to “Experimental procedures”.

| Compared parameter (statistical method) | Model | Sample size per dietary group or combination | Sex-specific sample size per group | Sample size per tank | Total sample size |
| --- | --- | --- | --- | --- | --- |
| SL 15 days (LME) | SL15 ~ diet + (1\|tank) | 90 | 90J | 30J | 180 |
| SL 30 days (LME) | SL30 ~ diet + (1\|tank) | 32 | 16M:16F | 4M:4F | 64 |
| SL 56 days (LME) | SL56 ~ diet + (1\|tank) | 32 | 16M:16F | 4M:4F | 64 |
| BM 15 days (LME) | BM15 ~ diet + (1\|tank) | 90 | 90J | 30J | 180 |
| BM 30 days (LME) | BM30 ~ diet + (1\|tank) | 32 | 16M:16F | 4M:4F | 64 |
| BM 56 days (LME) | BM56 ~ diet + (1\|tank) | 32 | 16M:16F | 4M:4F | 64 |
| SGR (Gaussian LM) | SGR ~ diet + sampling point | 7 | - | - | 14 |
| Fulton's condition (LME) | K ~ diet + sex + (1\|tank) | 32 | 16M:16F | 4M:4F | 64 |
| VFS (MPA) | VFS ~ diet + error (tank) | 32 | 16M:16F | 4M:4F | 64 |
| LM (LME) | LM ~ diet + BMdis + (1\|tank) | 32 | 16M:16F | 4M:4F | 64 |
| Additional LM (LME) | LM ~ diet + HVE + BMdis + (1\|tank) | 19-21 | 10M:10F | 2-3M:2-3F | 40 |
| HVE (LME) | HVE ~ diet + sex (1\|tank) | 20-21 | 10M:10-11F | 2-3M:2-3F | 41 |
| THCV | Pearson's $\chi^2$ test | 19-21 | 10M:10F | 2-3M:2-3F | 40 |
| N of eggs (Neg.Bin. GLMM) | N eggs ~ pair combination + BM + (1\|tank) | 16 | 16M:16F | 4M:8F | 32/2358 |
| FR (Bin. GLMM) | FR ~ pair combination + (1\| TM.TF) | 16 | 16M:16F | 4M:8F | 32/2358 |
| Ovary mass (LME) | OM ~ diet + BMdis + (1\|tank) | 16 | 16F | 4F | 32 |
| Egg diameter (LME) | ED ~ diet + (1\|tank/ID.female) | 16 | 16F | 4F | 32/837 |
| 30dpf egg survival (Bin. GLMM) | ES ~ diet + (1\|dish) | 100 | - | - | 400 |
| 30 dpf egg diapause stage | Pearson's $\chi^2$ test | 25-31 | - | - | 114 |

The dietary groups differed in four of the nine somatic parameters we measured (Fig. 1). Over 35 days of either the pellet or bloodworm diet (corresponding to the period of most intense growth) fish increased their body mass, on average, from 0.096 g to 1.203 g in females and 2.547 g in males (Fig. 1a).

**Figure 1:**
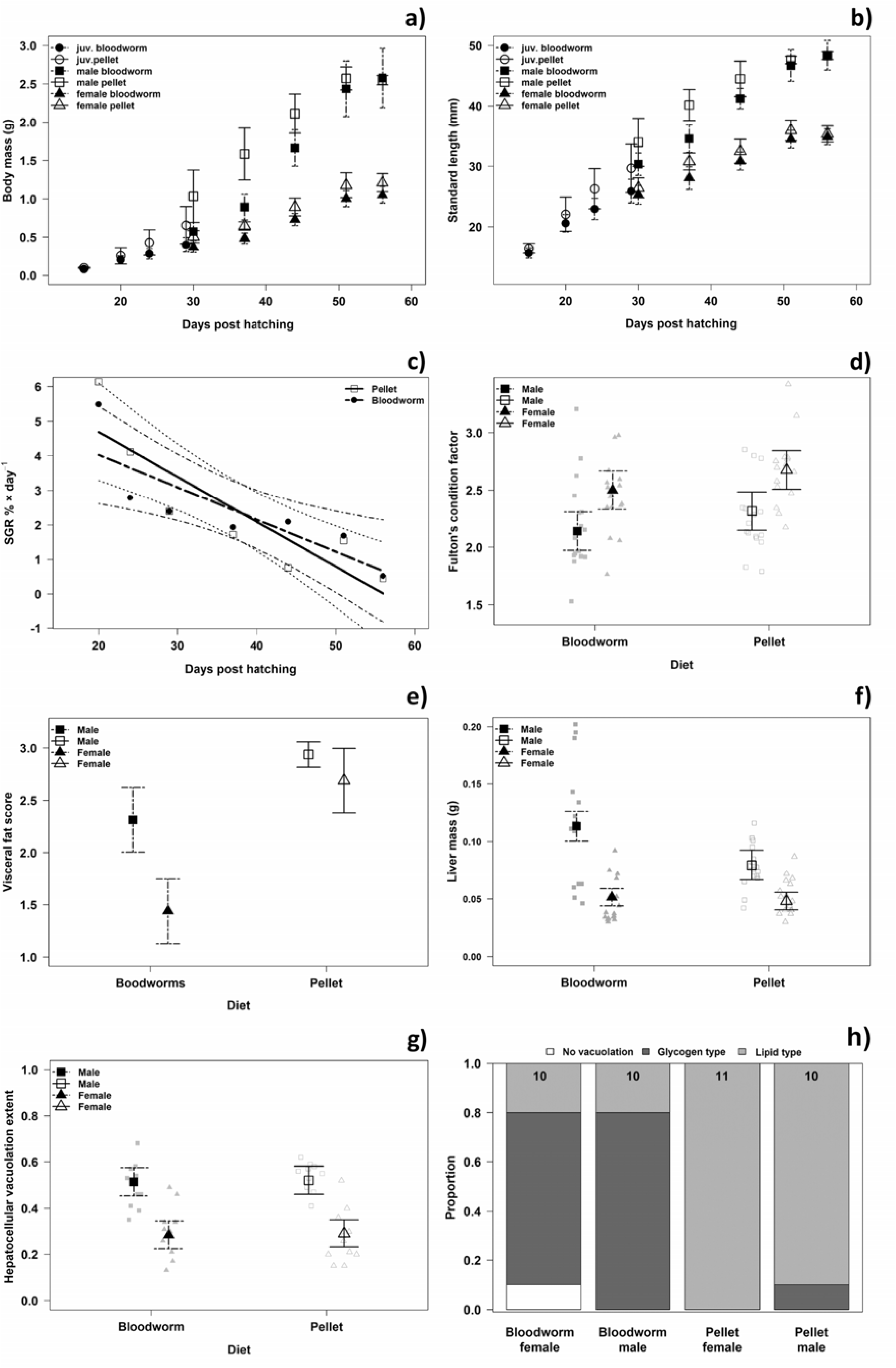
Somatic parameters in the two dietary groups. **a)** Age-dependent body mass. **b)** Age-dependent standard length. Values in a) and b) for juveniles are tank dependent weighted means (weighted by number of fish). Confidence intervals (CI, 95%) are computed from tank-specific weighted means. **c)** Specific growth rate of body size in % per day during whole experimental period in both diet groups. Linear model fit with 95% CI. SGR for body mass is not shown given its similarity with body mass SGR. **d)** Fulton’s condition factor with raw data points and model estimated means with 95 % CI. **e)** Visceral fat score. The values represent sex and tank dependent means and their associated CIs. **f)** Liver mass corrected for eviscerated body mass. Model estimated means with 95% CI and original data points. **g)** Relative hepatocyte cytoplasmic vacuolation. Model estimated means with 95% CI and original data points. **h)** Proportion of hepatocyte cytoplasmic vacuolation types computed from raw data. Numbers in upper parts of bars indicate sample sizes. Note that observation points in plots do not necessarily fit to the presented means due to the role of random factors. Figures without observation points were based on tank-specific means.

Individual body size did not differ between treatments (start of the experiment at 15 dph: Linear mixed effect model (LME), P = 0.196; end of juvenile period at 30 dph: males: P = 0.280, females: P = 0.287; end of the experiment at 56 dph, males: P = 0.973, females: P = 0.751, Fig 1b). Body mass was higher in the pellet diet treatment (age 15 dph: 19% difference, P = 0.003) and this difference increased considerably by the end of the juvenile period (age 30 dph: males: 80%, P = 0.012, females: 28%, P = 0.009, Fig. 1a). At the end of the experiment, there was no difference in male body mass (age 56 dph: P = 0.920) and only a minor difference in female body mass (15%; P = 0.017, Fig. 1a). The specific growth rate was similar in both groups (Linear regression, body size: P = 0.949, body mass: P = 0.960; Fig. 1c). Similar growth rates in both dietary treatments were accomplished by a higher consumption of bloodworms compared to pellets (Fig. S1). This resulted in a Food conversion ratio (FCR) of 7.44 (range 5.39-10.36) for bloodworms and 0.99 (0.67-1.93) for pellets.

Body mass differences between dietary treatments could be related to body condition but Fulton’s condition factor was similar between the treatments (7% difference, LME, P = 0.0503, Fig. 1c). Visceral fat score was higher in the pellet diet (males: 27%, Multistratum permutation analysis, P = 0.015; females, 87 %, P < 0.001, Fig. 1d). A sex-specific effect of diet on liver mass was detected. Liver mass (controlled for eviscerated body mass) did not depend on diet in females (7%, LME, P = 0.0531, Fig. 1e) while bloodworm-fed males had higher liver mass than pellet-fed males (42%, P < 0.001, Fig. 1e).

Histological examination of the liver parenchyma revealed two essential types of hepatocellular vacuolation (Fig. S3). (i) The lipid type manifested itself as unstained cytoplasmic vesicles with sharp edges. (ii) The glycogen-like type was characterized by irregular vacuoles containing slightly flocculent material. The presence of glycogen was evidenced by PAS (periodic acid Schiff reaction) positive staining of uneven sized grains dispersed in the cytoplasm (Fig. S3e). Fish fed pellets had almost exclusively lipid type hepatocellular vacuolation while fish fed bloodworms had mostly the glycogen-like type (χ^2^ test, P = 0.003, Fig. 1g, Fig. S3). Out of 41 histologically examined fish, only a single female (fed bloodworms) had no sign of hepatocellular vacuolation (Fig. 1 a). Hepatocellular vacuolation extent was similar in both treatments (LME, P = 0.838, Fig. 1f) but males were more affected than females (P < 0.001). Male livers with glycogen-like type vacuolation were larger than livers with lipid type vacuolation (54 %, LME, P < 0.001, Fig. S2) but male liver mass was independent of the extent of hepatocellular vacuolation (P = 0.461). In contrast, female liver mass was independent on vacuolation type (P = 0.260) but depended on vacuolation extent (P < 0.001, Fig. S2). Pre-neoplastic lesions were found only in one fish fed bloodworms and three fed pellets, too few for statistical comparison.

### Reproductive parameters are not compromised by pelleted diet

While fish reproductive parameters typically depend on diet (Moatt et al. 2019), none of the six reproductive parameters we measured differed between the treatments (Fig. 2). Therefore, egg number (Negative binomial Generalized linear mixed effect model (GLMM), P = 0.242, Fig. 2a), fertilization rate (Binomial GLMM, P = 0.203, Fig. 2b), ovary mass (controlled for body mass, LME, P = 0.624, Fig. 2c), egg size (LME, P = 0.927, Fig. 2d) or egg survival over 30 days post fertilization (30 dpf; Binomial GLMM, P = 0.814, Fig. 2e) were not compromised by the pellet diet. Note that the egg number was not different even for a contrast in female diet only (GLMM, P=0.114). In addition, there was no difference in the egg developmental stages at 30 dpf (Pearson’s χ^2^ test, P = 0.054, Fig. 2f), suggesting comparable effects of the two diets on embryo development of the next generation.

**Figure 2:**
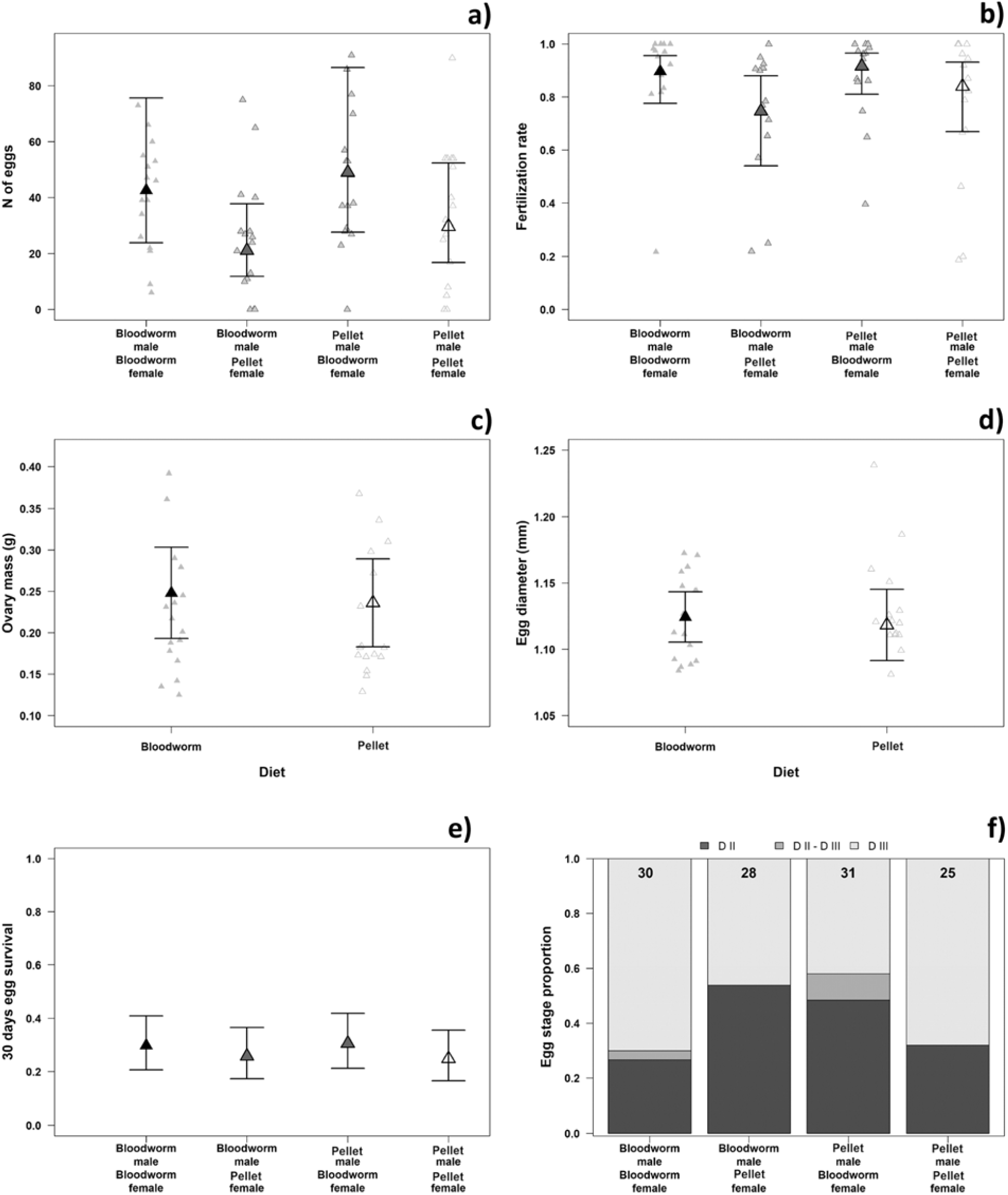
Reproductive parameters in pellet and bloodworm dietary groups. **a)** Fecundity. Raw data points and Negative-Binomial GLMM estimated means and 95% confidence intervals (CI) of four combinations of pairs. **b)** Proportion of fertilized eggs from four pair combinations. Means and 95% CI are Binomial GLMM estimated values. **c)** Reproductive allotment indicated by ovary mass corrected for eviscerated body mass. Model (LME) estimated means and 95% CI and raw observation points. **d)** Egg diameter. Model (LME) estimated mean and 95% CI. Observation points are female-specific mean egg sizes. **e)** 30 days post fertilization (dpf) survival of egg individually incubated in water. Mean and 95% CI are Binomial GLMM estimated values. Raw data are not presented due to their binomial nature. **f)** Egg diapause stage proportion after 30 dpf. Values in the upper part of bars are sample sizes from which diapause stage was determined.

## DISCUSSION

The standardized diet is necessary for full establishment of model organisms in aging. The present study compared Turquoise killifish key fitness traits between two diets – widely used bloodworms (Polačik et al. 2016) and BioMar INICIO formulated pellets. Overall acceptance of pellets was good from puberty (21 days old) onwards. A minor dietary effect was found on body mass probably via differences in visceral fat storage. Differences in diet-dependent energy storage also occurred at the hepatocellular level. A sex-specific effect of diet was detected in liver mass with heavier livers in bloodworm-fed males but not females. Standard length, Fulton’s condition, extent of hepatocellular vacuolation and reproductive parameters did not differ significantly between diets suggesting the suitability of the pelleted diet as a standard for *N. furzeri*. The similar performance of bloodworm-fed fish compared to pellets-fed fish was compensated by 7.5-fold higher mass of consumed bloodworms. We believe that this study represents a promising demonstration of establishing a standardized formulated diet as a prerequisite for inter-laboratory comparison between studies using *Nothobranchius furzeri*.

Growth rate often affects lifespan (Polačik et al. 2014; Vrtílek, Žák, Pšenička, et al. 2018) and, at the same time, it is a widely used parameter to compare diet quality (Watanabe et al. 1991; Goolish et al. 1999). Turquoise killifish body size was similar for both diets during entire experimental period. Body mass (but not body size) was initially slightly higher in pellet-fed fish but the difference disappeared in males upon reaching their asymptotic size (56 days old). Initial higher body mass in the pellet group can be related to the marginally higher proportion of males in that group (Table S1), but the sex of juveniles is indistinguishable and the male-bias was only detected after fish reached maturity. Body mass also increased faster in the pellet-fed group than in the bloodworm-fed group between age 15 and 30 days. This can be associated with faster growth of pellet males compared to bloodworm males and consequently their earlier maturation related to faster development of sexual characteristics such as deeper body and development of gonads. In addition, the higher fat deposition could have already been manifested at that age. Altogether the results suggest better nutritional status in pellet diet group than bloodworm group, likely related to the higher lipid content of pellets.

Condition is often used as a marker of health status in fish (Moatt et al. 2019). Fulton’s condition factor did not differ between dietary groups, but the visceral fat score in the pellet group was much higher, indicating their superior somatic condition. This inconsistency can be related to significantly bigger livers in bloodworm-fed males which could mask the tendency for lower condition in bloodworm-fed group. All fish were fasted prior to dissection and had empty guts, hence, differential gut fullness could not have been responsible for the difference in condition factor. High visceral fat loads in fish fed on pellets are common (Penrith et al. 1994; Moatt et al. 2017) but this does not necessarily shorten lifespan (Moatt et al. 2019). The most probable cause of high visceral fat in pellet-fed fish is that the fat content in pellets is 3-5 times higher than bloodworms (Table 1). The increased level of fat accumulation could potentially become a study model for dietary induced obesity (Oka et al. 2010). At the same time, high fat accumulation suggests that formulation of a Turquoise killifish specific formula for pellets is desirable.

Fish hepatocytes are more vacuolated than mammal hepatocytes (Wolf & Wolfe 2005). Wild annual killifish have an exceptionally high extent of hepatocellular vacuolation (Vrtílek, Žák, Blažek, et al. 2018; Godoy et al. 2019) suggesting that it is a natural state. In the present study, male livers were more vacuolated than females’ which is in accordance with previous findings both in captivity (Di Cicco et al. 2011) and the wild (Vrtílek, Žák, Blažek, et al. 2018). Fish fed pellets possessed almost exclusively lipid type hepatocellular vacuolation which is in accordance with other studies of fish fed on pellets (Penrith et al. 1994; Wolf & Wolfe 2005). On the other hand, the bloodworm group had a similar extent of hepatocellular vacuolation, though predominantly of the glycogen-like type. Higher liver glycogen reserves develop when fish consume a diet rich in carbohydrates (Kohla et al. 1992) and bloodworms had 2-3 times higher carbohydrate content than pellets. This differentiation between the two types of energy reserves in livers provides an interesting model for comparative physiology.

The sex-specific reproductive role affects liver size (Casselman & Schulte-Hostedde 2004). Overall, males had larger livers than females and female liver size did not differ between dietary groups. We speculate that the lack of difference in female liver size between dietary treatments is related to energetically demanding oocyte production, directly linked to liver metabolism (Papa et al. 2019). Higher reproductive energy mobilization in females is also indicated by their lower hepatocellular vacuolation extent and lower visceral fat accumulation. Male gamete production is less energy demanding (Trivers 1972) and allows them to store more energy reserves in their livers. Glycogen storage increases liver size (Hemre et al. 2002) and extensive glycogen reserves were found in bloodworm-fed males, making this the most likely cause of their large livers.

The dominance of glycogen-like vacuolation in hepatocytes of the males kept on bloodworm diet accentuates the need to distinguish between physiological and pathological accumulation of glycogen. The solution lies in the understanding subcellular changes, which requires information from transmission electron microscopy. Similarly, simultaneous occurrence of PAS positive large vacuoles and nuclei in submembrane position deserves further attention. Our findings seem to contradict the widely accepted histological definitions of glycogen vacuolation in fish by Wolf & Wolf (2005). For our experimental fish we suggest a two-step cell alteration, i.e., lipid macrovesicular vacuolation followed by glycogen accumulation.

Generally in fish, a pellet diet affects the reproductive parameters of females (Markovich et al. 2007; Mandal et al. 2012) but less so of males (Henrotte et al. 2010; Moatt et al. 2019). The present study partly supports this, because there was no difference in fertilization ability in pellet fed males but a tendency for lower fecundity and fertilization rate in females. This suggests that female lifespan may be affected via reproduction-body maintenance trade-off (Shanley & Kirkwood 2000). Otherwise we did not detect any effect of diet on female reproductive allocation (ovary mass and egg size) indicating that females rather respond to feeding periodicity (Vrtílek & Reichard 2015) than to macronutrient composition. There was no difference in the developmental stage at 30 dpf, suggesting comparable effects of two parental diets on the dynamics of embryo development.

Good acceptance and ingestion of a newly introduced diet is necessary for its successful establishment in laboratory studies. In the present study, BioMar INICIO - commercially available formulated diet, was well accepted by killifish from puberty onwards. Pellets were better ingested by fish (indicated by lower FCR) and consequently fish produced fewer feces, reducing the need of frequent water exchange. The challenging transition of juvenile fish to a formulated diet is a common problem in aquaculture (Goolish et al. 1999). It is possible to feed killifish pathogen free, but nutritionally unstable, bloodworms during the juvenile transition period to pellets (e.g. Hikari Bio-Pure bloodworms, Japan, http://www.hikariusa.com). However, we have a recent experience that transition from *Artemia* nauplii directly to pellets is possible (Žák, personal observation). Yet, in Turquoise killifish, it is necessary to train fish for satisfactory pellet acceptance. We highlight that the age around puberty and maintenance in social tanks are optimal conditions for a successful killifish transition to pellets. We do not believe that the short period when killifish were co-fed bloodworms and pellets affected our results, because fish were fed solely on pellets for 35 days when they gained 4-10 times their initial body mass. However, we acknowledge that the co-feeding transition period could be a time window for disease or chemical contamination from bloodworms (Papa et al. 2019). Otherwise, the combined diet of commercial pellets with live food improves the reproductive parameters of fish (Mandal et al. 2012; Vasagam et al. 2007).

The present study demonstrates that Turquoise killifish can be kept on a diet of dry pellets, potentially enabling research into the effect of macronutrient manipulation on life histories (Moatt et al. 2019). We believe that the optimal formulated diet should contain a lower proportion of fat than that contained in the pellets used in the present study and a lower proportion of carbohydrates than bloodworms. We believe that our findings are applicable to most medium to large-sized *Nothobranchius* species; we were also able to convert adults of *N. orthonotus, N. kadleci* and *N. guentheri* to a formulated diet (Žák, personal observation). In general, feeding pellets is more practical than bloodworms and it is less financially demanding. We recommend that a standardized formulated diet for *N. furzeri* should be universally adopted.

## EXPERIMENTAL PROCEDURES

### Killifish origin and housing

All experimental work was completed on the wild-derived strain MZCS 222 (Cellerino et al. 2016) of Turquoise killifish (*Nothobranchius furzeri*). Detailed husbandry and size assortment is described in Supplementary material (Killifish housing, Table S1). In short, Fish were hatched and raised in the common tanks until the age of 12 days following (Polačik et al. 2016). Thereafter they were moved to 35L tanks with three replicates (cca 30 fish per tank) per dietary treatment and size assorted at 15, 17 and 20 days (Table S1) from the initial density of 30 fish per tank (n = 180 fish, 3 tanks per treatment) to 10 – 21 fish per tank (5 tanks per treatment). Size assortment was done to improve the growth and to reduce aggression (Polačik et al. 2016). Keeping fish in social groups improves fish willingness to feed and therefore promotes easier recognition of new fooditems (Lepič et al. 2017). At the age of 29 days, the final experimental groups (four replicates per treatment, 4 males + 8 females per 35L tank) were established. The experiment was terminated when fish reached asymptotic growth at the age of 56 days. Water temperature was kept at 27.1 °C ± 0.87 (mean ± SD) and light regimen was 14L:10D.

### Killifish feeding procedure

After hatching, the fish were fed live *Artemia* nauplii (Sanders, USA, www.gsla.us) three times per day (8:30, 13:15, 18:15) in order to provide continuous access to live *Artemia*. At present, live food is irreplaceable at the early juvenile stage of *Nothobranchius* spp. Finely chopped bloodworms (Chironomidae, Petr Grýgera, Czech Republic, https://nakrmryby.cz/) started to be added to the diet at the age of 12 days. From this point onward, the control group was fed exclusively with bloodworms but in the experimental group we started to supplement the bloodworms with the finest grade of dry food pellets BioMar INICIO 0.4 mm (Denmark, https://www.biomar.com/). At the age of 15 days, several fish were observed to accept pellets. Pellets were soaked in water for 1-3 min before to soften them before being added to the tank in batches of 1 to 5 pellets using a Pasteur pipette. Fish were fed *ad libitum* (amount consumed within 5 min, Fig. S1) feeding with bloodworms or pellets three times per day. The pellet size was gradually increased in 0.2 mm steps (from 0.4 mm to 1 mm and 1.5 mm; Figure S4), larger pellets being offered prior to the smaller ones until they were fully adapted to the larger size (usually 2-3 days). Detailed information on pellet size with respect to the experimental stage can be found in Supplementary Fig. S4. The mixed diet (bloodworms and pellets) continued up to the age of 21 days, when all experimental fish fully accepted the pellets. At this point the feeding schedule was reduced to twice a day (11:30, 18:45). The age of 21 days coincided with the onset of coloring up in males. From this point onwards, experimental fish received exclusively pellets until the termination of the experiment. Dry pellets were fed from the age of 38 days and were vigorously accepted.

Prior to each feeding, food mass was determined. Before feeding, thawed bloodworms were left for 5 minutes in the sieve to dry and then weighed to the nearest 0.001 g using an analytical scale (Kern PCB 350-3, www.kern-sohn.com, Germany). The same procedure was done with unused bloodworm after feeding. The difference in mass before and after feeding was taken as the amount of bloodworms consumed. A smaller dose of pellets than the fish were expected to consume was weighed to the nearest 0.001 g in small plastic cup prior each feeding. Then small weighted amounts of pellets were given to fish until they were fully satiated.

The feed conversion ratio (FCR = food intake / body mass gain) was computed for each diet at each body mass sampling point. The macronutrient content (Table 1) of bloodworms (and BioMar INICIO 0.8 mm as a control for method calibration), were analyzed in an accredited metrologic laboratory at the National Veterinary Institute in Olomouc, Czech Republic (https://www.svuolomouc.cz/).

### Somatic parameters

We measured body size (SL, standard length: excluding caudal fin, mm), body mass (BM, g), specific growth rate (SGR, % × day^-1^), Fulton’s condition factor (K ; body mass divided by cubic SL and multiplied by 100), visceral fat score (VFS), liver mass (LM, g), hepatocellular vacuolation extent (HVE) and type of hepatocellular vacuolation (THCV) were determined. SL and BM were measured at five day intervals between age 15 and 30 days and at seven day intervals from 30 days to 51 days. The last measurement was completed at the age of 56 days. BM was measured on slightly wet towel-dried live specimens to the nearest 0.001 g using analytical scales. SL was measured using ImageJ (NIH, Bethesda, MD, USA) from photographs in a plastic container with shallow water and scale. SGR was computed from sampling-point-pooled averages for both SL and BM with the formula: SGR = (ln*S*_*2*_ – ln*S*_*1*_)×(100/*t*) where *S*_*2*_ is average terminal size (SL, mm or BM, g), *S*_*1*_ is initial average size and *t* is the length of time interval in days. K (Ricker 1975) was calculated at the end of the experiment. VFS was visually determined after opening the body cavity at the end of the experiment (scale 0 – 3; 0 – no visceral fat; 3 – internal organs completely covered by fat). LM of freshly extracted livers was weighted to nearest 0.001 g using analytical scales.

### Liver histology

The liver was removed from the body cavity, fixed in Davidson’s fixative (for 48 h), stored in 70 % ethanol (4 days) and processed using the paraffin technique. This included dehydration in series of ethanols, aceton and xylen, and embedding in Histoplast (Serva, Germany, https://www.serva.de/enDE). To enure the best possible prerequisites for comparison, liver samples taken from 41 individuals (10-11 per treatment-sex combination, 2-3 per sex-tank combination) were oriented identically while embedded into Histoplast blocks. From the parietal part of the liver, five semi-serial 4 µm sections per fish were prepared using a rotating microtome HM360. Of these, three sections per fish were stained with Mayer’s hematoxylin and eosin. The histological findings were documented using a BX60 Olympus microscope equipped with a DP71 camera under 175× magnification. The images of representative sections were analysed for HVE in ImageJ that calculated the proportion of total unstained area (mainly vacuoles) against the well-stained tissue. This method was validated by re-examining a subsample of 24 sections by another experienced evaluator and there was a strong association between both results (Pearson’s correlation coefficient, r = 0.76, t_22_ = 5.51, P < 0.001). THCV was evaluated blindly, based on vacuole character; glycogen-like type vacuolation (Fig. S3 c), e), g): irregular vacuoles with indistinct margins (Wolf et al. 2015); lipid type vacuolation (Fig. S3 b),d),f): a clear round vacuoles with sharp edges (Wolf et al. 2015). Several sections with liver tissue containing irregular vacuoles in the cytoplasm of hepatocytes (glycogen-like THCV) were validated by supplementary staining for glycogen (periodic acid Schiff reaction, PAS, Fig. S3 e).

### Reproductive parameters

To compare fecundity and fertilization rate between treatments, 16 males (from a total of four tanks) and 16 females (from a total of two tanks) per treatment were spawned in a 2L plastic container with substrate of 0.5 cm fine grained sand for 2 hours (Polačik et al. 2016) Experimental spawning took place at the age of 53 and 54 days respectively. Each female was spawned twice – in a random order with a male from the same treatment group and with a male from the other treatment group. This design was selected to address potentially confounding parental effect on reproductive parameters. Fish were spawned in this setting twice before (age 51 and 52 days respectively) for habituation and standardization. After experimental spawning, each female was weighed to 0.001 g and males and females were released into separate tanks to prevent spawning.

From each pair combination, survival of 100 individually incubated eggs (4 plastic dishes with 25 compartments) was determined. The maximum egg contribution from each female in a respective day to the egg pool from which 100 eggs were selected was 20. Eggs were incubated in 25°C water in a laboratory incubator (Q-Cell, Pol-lab, www.poll.pl). After 30 dpf, the developmental stage of the surviving eggs was determined (Furness et al. 2015). Reproductive allotment was estimated from ovary mass (Vrtílek & Reichard 2015) at the age of 56 days. Ovaries were extracted from the body cavity and fixed in 4% formalin for later measurement and egg extraction for egg size measurement (Vrtílek & Reichard 2015). Egg size was measured for a batch of 10 – 51 eggs per female.

### Statistical analysis

All statistical analysis was carried out in R environment 3.5.2 (R Core Team 2018). All possible two way interactions were included in models but removed when non-significant. In case of a significant interaction with sex (SL30; SL56; BM30; BM56; LM, VFS), separate sex-specific analyses were performed. For statistical analysis of SL30; SL56; BM30; BM56; LM, VFS and ovary mass, 4 females per tank were randomly selected to retain a balanced dataset. Statistical difference of K was interpreted only between dietary groups but not between sexes, as the sex difference is not informative in sexually dimorphic species (Peig & Green 2010). In THCV statistical analysis, a single individual with no hepatocyte vacuolation was omitted due to its rarity. Additional analysis revealing the effect of HVE on liver size was completed with THCV as a fixed factor (instead of treatment as the fixed factor) due to their strong collinearity (Fig. 1g). The proportion of individuals with pre-neoplastic lesions was not too few to analyze (4 individuals out of 41 examined). A detailed overview of sample sizes and statistical models can be found in Table 1. Complete tables of statistical results can be found in the supplementary excel sheet (Supplementary: Statistical_results.xlsx). Original datasets (Žák et al. 2019) are available via Figshare repository (10.6084/m9.figshare.9825068).

## Supporting information

All Supplementary Material

## ACKNOWLEDGEMENTS

We thank Václav Homolka and Jiří Farkač for their help with killifish maintenance, to Gabriela Vágnerová for her help with histological slide preparation and to Matej Polačik, Milan Vrtílek and Radim Blažek for insightful comments on the manuscript. We thank Rowena Spence for English correction. Funding came from the Czech Science Foundation (19-01781S). We declare no competing interest. Original data are available via Figshare repository (doi: 10.6084/m9.figshare.9825068).

## AUTHOR CONTRIBUTIONS

JŽ designed the study

ID did the histological analysis

JŽ and ID collected the data

JŽ analysed the data

JŽ, MR and ID wrote the manuscript

