## Supplementary material for "Performance of Turquoise killifish, model organism in aging, on commercial pelleted diet: a step towards husbandry standardization": All Supplementary Material

KILLIFISH HOUSING

All experimental work was completed on the wild-derived strain MZCS 222 (Cellerino et al. 2016) of Turquoise killifish (*Nothobranchius furzeri*) bred at the accredited breeding facility at the Institute of Vertebrate Biology from 26 March to 21 May 2019. Fish were hatched and raised in common tanks until the age of 12 days following (Polačik et al. 2016). At the age of 12 days, fish were moved to six 35L glass tanks (30 fish per tank, three replicates for the bloodworm treatment and three replicates for pellets) with an air-powered sponge filter. The light regime was 14L:10D. Water temperature was maintained at 27.1 °C ± 0.87 (mean ± SD, recorded at 1 hr interval by two HOBO UA-002–64 loggers, Onset Computer, Bourne, MA, USA). Fish were sorted by size at the age of 15, 17 and 20 days to reduce the negative effects of aggression and fish density on growth (Polačik et al. 2016; Vrtílek et al. 2019; Table S1). At the age of 29 days, final experimental groups (four replicates per treatment, 4 males + 8 females per 35L tank) were established. A female-biased sex ratio was chosen to reduce female harassment and male-male aggression (Polačik et al. 2016). Keeping fish in social groups improves fish willingness to feed and therefore promotes easier recognition of new food items (Lepič et al. 2017). Any dead or belly-sliding (common in captive *Nothobranchius* spp. (Dyková et al. under Review)) fish were replaced by a similarly-sized individual from an additional tank with an identical setup to the experimental tanks. The number of replaced fish between dietary groups was similar (12 in the bloodworm treatment and 8 in the pellet treatment). The experiment was terminated when fish reached asymptotic growth at the age of 56 days.

SUPPLEMENTARY FIGURES

FIGURE S1

Figure S1: Mean food mass consumed by one killifish during one feeding event. Values are the mean from all feeding events on a given day. Note the different y-axis scale for each diet. Dashed line is age when all fish were adapted to pellets. a) pellets, b) bloodworms

FIGURE S2

Figure S2: Relationship between liver mass and extent of hepatocellular vacuolation. Liver mass in females increases with the extent of hepatocellular vacuolation which was not observed in males. Points of observation represent raw data. Lines are simple trends from linear relationships.

FIGURE S3

Figure S3: Various types of hepatocellular vacuolation of *Nothobranchius furzeri* and extent of hepatocellular vacuolation stained with Mayer’s hematoxylin and eosin (a), b), c), d), f), g) and periodic acid Schiff reaction (e) under 175× magnification. a) Liver parenchyma with no apparent hepatocellular vacuolation, b) macrovesicular lipid type hepatocellular vacuolation, in analysis considered as lipid type vacuolation, c) glycogen-like type severe hepatocellular vacuolation, in analysis considered as glycogen-like type vacuolation, d) mixed size of lipid vacuoles, in analysis considered as lipid type vacuolation, e) Section with glycogen-like type vacoulation stained with periodic acid Schiff reaction. Note stained content of vacuoles. f) mostly microvesicular lipid type vacuolation, in analysis considered as lipid type of vacuolation, g) glycogen-like type of minimal hepatocellular vacuolation, in analysis considered as glycogen-like type vacuolation.

FIGURE S4

Figure S4: Schematic description of feeding regime (formulated diet BioMar INICIO) and killifish husbandry. Black cell stands for activity performed. See chapters “Killifish origin and husbandry” and “Killifish feeding procedures” for details.

Figure S1

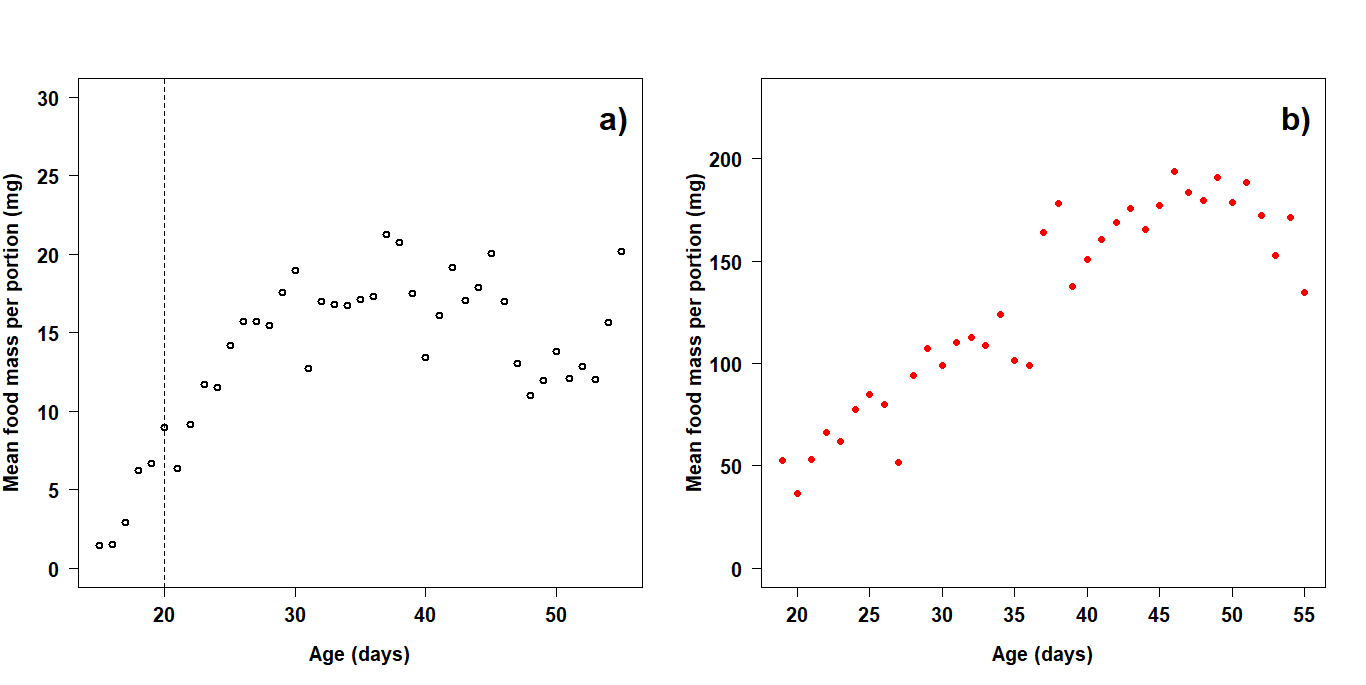

Figure S2

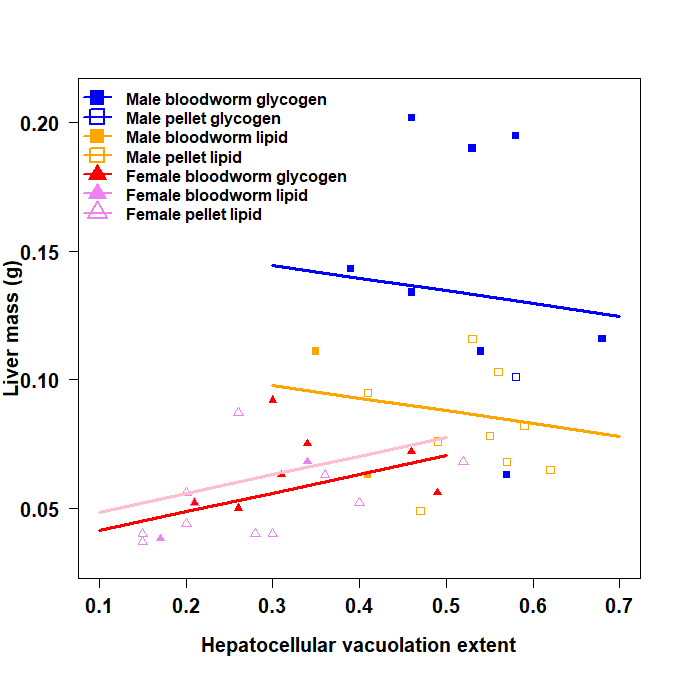

Figure S3

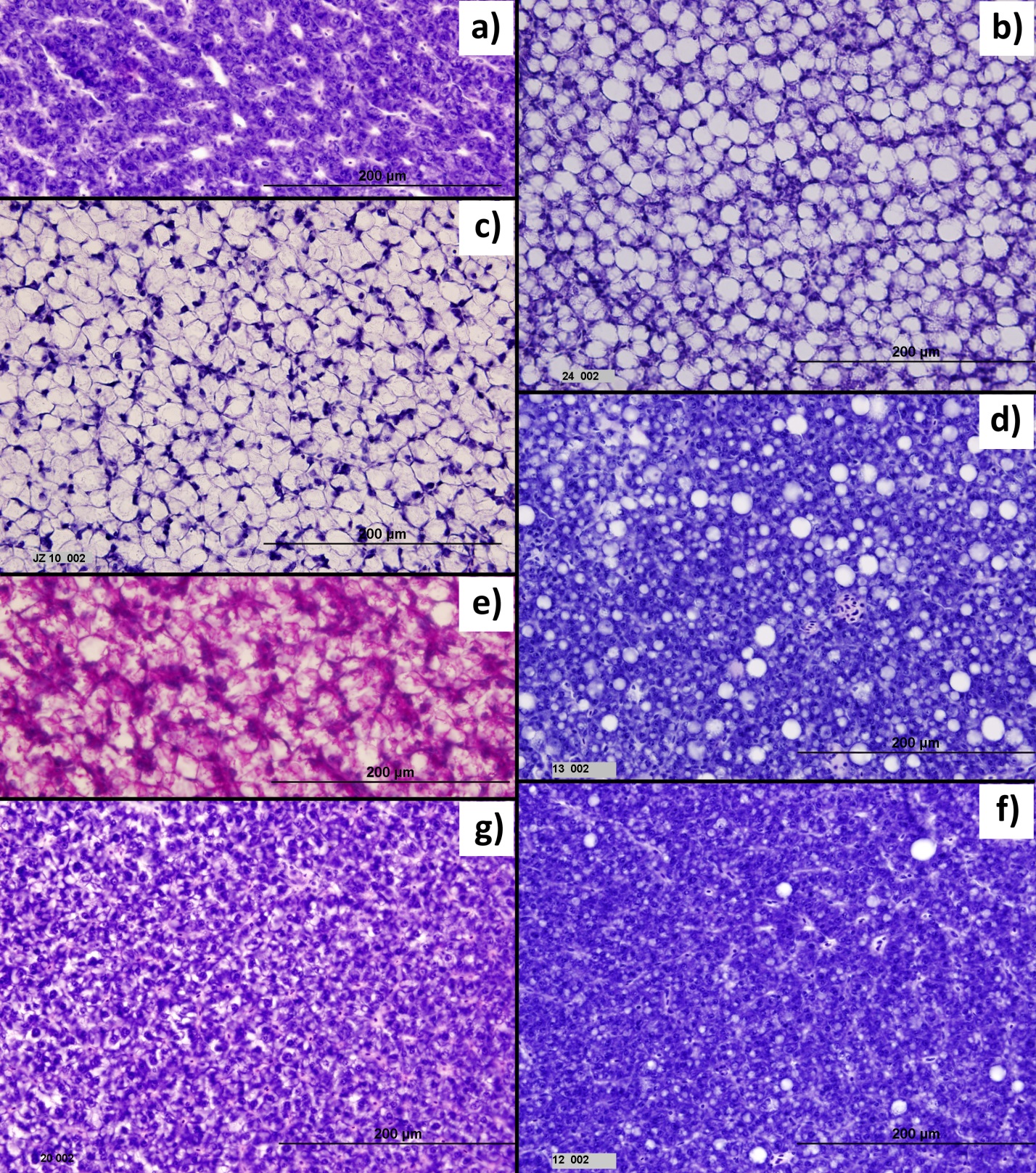

Supplementary Figure S4

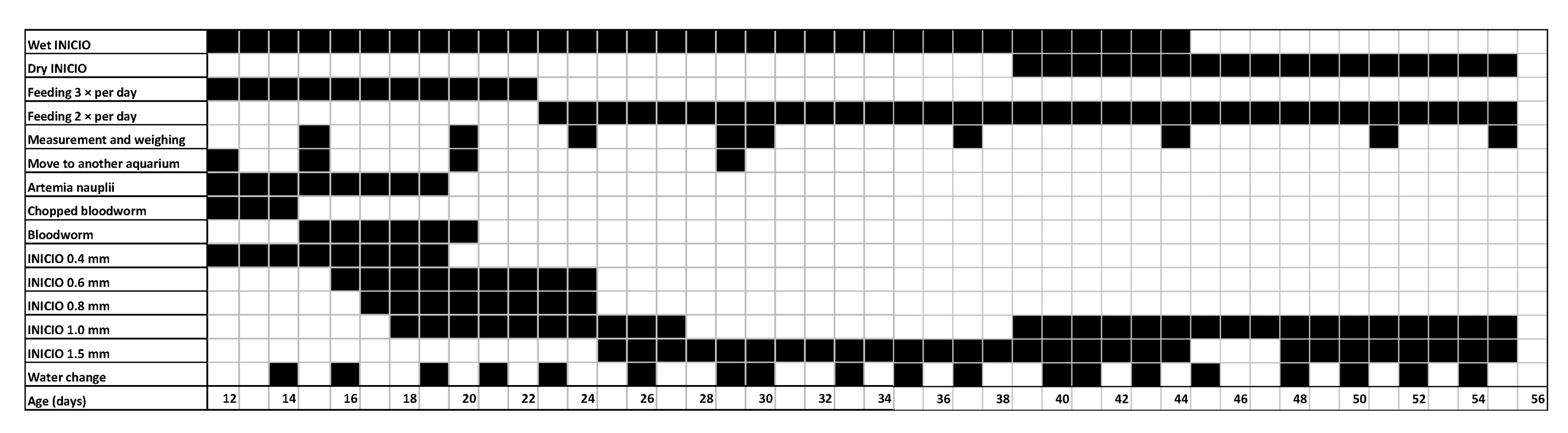

.

SUPPLEMENTARY TABLES

TABLE S1

Table S1: Overview of fish density during the experiment. During the juvenile period regular size assortment of fish was needed to achieve balanced growth. INICIO stands for BioMar starter food INICIO and number behind is for the pellet size fed to killifish. Experimental group “B” stands for bloodworm dietary group and “F” stands for formulated diet. Sex ratio was determined at the end of juvenile period before assembling final experimental tanks. Killifish size group is a result of regular fish assortment; similarly sized individuals were released together to have equal chance to acquire food. Small: the smallest individuals, Average: medium sized individuals, Large: largest individuals.

| Age (days) | Tank volume (L) | Number of tanks | Experimental group | Diet | Number of fish per tank | Sex ratio  M:F | Fish size group |
| --- | --- | --- | --- | --- | --- | --- | --- |
| 0-3 | 6 | 1 | - | *Artemia naupli* | Cca 200 | J | All |
| 3-12 | 75 | 2 | - | *Artemia naupli* | Cca 90 | J | All |
| 13-14 | 35 | 3 | B | Chopped bloodworms + *Artemia naupli* | 30 | J | All |
| 13-14 | 35 | 3 | F | Chopped bloodworms + *Artemia naupli* | 30 | J | All |
| 15-16 | 35 | 3 | B | Bloodworms + *Artemia naupli* | 25 | J | Average |
| 15-16 | 35 | 3 | F | Bloodworms + INICIO 0.4 + *Artemia naupli* | 25 | J | Average |
| 15-16 | 35 | 1 | B | Bloodworms | 15 | J | Large |
| 15-16 | 35 | 1 | F | Bloodworms + INICIO 0.4 | 15 | J | Large |
| 17-19 | 35 | 1 | B | Bloodworms + *Artemia naupli* | 10 | J | Small |
| 17-19 | 35 | 1 | F | INICIO 0.4 + Bloodworms + *Artemia naupli* | 11 | J | Small |
| 17-19 | 35 | 1 | B | Bloodworms | 15 | J | Large |
| 17-19 | 35 | 1 | F | INICIO 0.8 +INICIO 1 | 15 | J | Large |
| 17-19 | 35 | 3 | B | Bloodworms | 20-21 | J | Average |
| 17-19 | 35 | 3 | F | Bloodworms + INICIO 0.4 + INICIO 0.6 | 20-21 | J | Average |
| 20-28 | 35 | 3 | B | Bloodworms | 15-17 | 14:32 | Average |
| 20-28 | 35 | 3 | F | INICIO 0.8 + INICIO 1 | 15-17 | 18:30 | Average |
| 20-28 | 35 | 2 | B | Bloodworms | 11 & 14 | 20:5 | Large |
| 20-28 | 35 | 2 | F | INICIO 1 | 12 & 16 | 25:3 | Large |
| 20-28 | 35 | 1 | B | Bloodworms | 9 | 7:2 | Small |
| 20-28 | 35 | 1 | F | INICIO 0.6 | 10 | 6:4 | Small |
| 29-56 | 35 | 4 | B | Bloodworms | 12 | 4:8 | All |
| 29-56 | 35 | 4 | F | INICIO 1 + INICIO 1.5 | 12 | 4:8 | All |

TABLE S1
